# Increased thermal tolerance under anoxic conditions in an extremophile fish from hot sulfur springs in Mexico

**DOI:** 10.1101/2023.07.21.550037

**Authors:** Korbinian Pacher, Natalia Hernández-Román, Alejandro Juarez-Lopez, Jesús Emmanuel Jiménez-Jiménez, Juliane Lukas, Yunus Sevinchan, Jens Krause, Lenin Arias-Rodríguez, David Bierbach

**Affiliations:** Department of Biology and Ecology of Fishes, Leibniz-Institute of Freshwater Ecology and Inland Fisheries, 12487 Berlin, Germany; Faculty of Life Sciences, Albrecht Daniel Thaer-Institute, Humboldt University of Berlin, 10115 Berlin, Germany; División Académica de Ciencias Biológicas, Universidad Juárez Autónoma Tabasco, 86150 Villahermosa, Mexico; Excellence Cluster Science of Intelligence, Technische Universität Berlin, 10587 Berlin, Germany; Institute for Theoretical Biology, Humboldt-Universität zu Berlin, 10115 Berlin, Germany; Bernstein Center for Computational Neuroscience, Berlin, 10099 Berlin, Germany

**Author notes:** Corresponding author’s. equal contribution / shared first authorship (Korbinian Pacher, Natalia Hernández-Román). equal contribution / shared senior authorship (Lenin Arias-Rodríguez, David Bierbach).

## Abstract

The thermal ecology of ectotherm animals has gained considerable attention in the face of human induced climate change. Particularly in aquatic species the experimental assessment of critical thermal limits (CT_min_ and CT_max_) may help to predict possible effects of global warming on habitat suitability and ultimately species survival. Here we present data on the thermal limits of two endemic and endangered extremophile fish species, inhabiting a geothermally-heated and sulfur-rich spring system in Southern Mexico: The sulfur molly (*Poecilia sulphuraria*) and the widemouth gambusia (*Gambusia eurystoma*). Besides physiological challenges induced by toxic hydrogen sulfide and related severe hypoxia during the day, water temperatures have been previously reported to exceed those of nearby clearwater streams. We now present temperature data for various locations and years in the sulphur spring complex and conducted laboratory thermal tolerance tests (CT_min_ and CT_max_) both under normoxic as well as sever hypoxic conditions in both species. Average CT_max_ limits did not differ between species under normoxic conditions. Surprisingly *P. sulphuraria* was found to reach a higher critical temperature (CT_max_ = 43.2°C) when tested under hypoxic conditions, while *G. eurystoma* on average had a lower CT_max_ when oxygen was absent. Based on this data we calculated both species’ thermal safety margins and used a TDT (thermal death time) model framework to relate our experimental data to observed temperatures in the natural habitat. Our findings suggest, that both species live near their thermal limits during the annual dry season and are locally already exposed to temperatures above their critical thermal limits. We discuss these findings in the light of possible physiological adaptions of the sulfur-adapted fish species and the anthropogenic threats for this unique system.

## Introduction

Anthropogenic global warming may lead to seasonal temperature increases by up to 4°C by 2100, along with an increase in the frequency of localized acute and extreme warming events (Comte and Olden 2017, Collins et al. 2019, Bierbach et al. 2022). These changes are likely to cause population declines, or even extinction of species that are poorly adapted to cope with these novel environmental conditions (Pacifici et al. 2015, Dai et al. 2022). A key trait and a critical factor for survival in this regard is an organism’s ability to survive and function within a specific temperature range, which is referred to as thermal tolerance. In its broadest sense, it can be defined by a species’ lower and upper physiological thermal maxima (e.g., critical thermal limits, CT_min_ and CT_max_, (Beitinger and Lutterschmidt 2011)), although growth and reproduction may require a narrower range thus higher minima and lower maxima (Pörtner 2001, Burleson and Silva 2011b, Illing et al. 2020). Understanding how organisms deal with increased or even extreme temperature regimes that will become more frequent becomes crucial for predicting the effects of climate change on populations and species, and ultimately may help to take suitable measures and actions to counteract local population declines or extinction events (Dai et al. 2022, Earhart et al. 2022, Desforges et al. 2023).

To this end, organisms that inhabit extreme thermal habitats may offer a unique opportunity to gain insights into the evolution of possible adaptations that allow survival in their extreme environments (Hillyard 2011, O’Gorman et al. 2014, Plath et al. 2015) while those extremophiles are furthermore most vulnerable to only small shifts in their thermal environment as they can already be assumed to live at the edge of their physiological limits and are often also endemic with very localized distribution ranges and small population sizes (Plath et al. 2015, Tobler et al. 2018). Thermally-influenced aquatic habitats like hot springs are among the most localized extreme habitats that often harbor unique species compositions (O’Gorman et al. 2014). For example, the Devil’s hole in northern Mojave Desert harbors the only known wild population of the Devil’s hole pupfish (*Cyprinodon diabolis*) with an estimated population size of only a few hundred individuals (Tian et al. 2022). In summer the water temperature can rise up to 39°C in that partly cavernous spring complex (Gustafson and Deacon 1998) and adults have been found to be able to tolerate temperatures up to 44°C (CT_max_, (Hillyard 2011)). Other systems with extreme thermal regimes include temporal ponds and pools along tropical and subtropical floodplains that are often shallow and thus get heated up due to sun radiation (Hauber et al. 2011, Jung et al. 2020). But even in the colder parts of the hemispheres, systems like the geothermal Hengill spring complex in Iceland are suitable ground for several highly adapted thermophile organisms (O’Gorman et al. 2016).

Aquatic habitats that are permanently or temporarily heated often show degrees of environmental hypoxia, which is a reduction or lack of dissolved oxygen (Diaz 2001). While the oxygen solubility in water decreases with increasing temperatures also microbial and other oxygen-consuming processes are faster under higher temperature thus further depleting oxygen from the water (Diaz 2001, Earhart et al. 2022). In addition to these environmental interactions, high temperature and hypoxia also have interacting effects on exothermal organisms themselves. These interactions are thought to be mediated through the joint impacts of high temperature and hypoxia on metabolism (Pörtner 2001, Pörtner et al. 2017, Earhart et al. 2022). Rising temperatures increase the rates of chemical and biochemical reactions; and these thermodynamic effects result in increases in metabolic demand, which must be met with increases in metabolic energy supply for an organism to maintain energy balance. For many animals, this energy will be provided through aerobic metabolism, and can become limited when environmental oxygen declines (Schulte 2015). These effects at the biochemical level cascade up to affect processes across levels of biological organization, with profound effects on complex physiological processes, such as cardiovascular function, muscle contraction, metabolism, energy budgets, which impact organismal growth and performance as well as thermal and hypoxia tolerance (McBryan et al. 2013, Little et al. 2020, Ern et al. 2023). Since chronic extreme conditions as well as periodic extreme thermal events are predicted to become more common also in yet normal habitats, the combination of high temperature and hypoxia in the environment can become particularly devastating as reports of heat-related mass die-offs of fishes and other higher aquatic organisms are further increasing in frequency (Dai et al. 2022). For example, the mass mortality events known as ‘summerkill’ that occur in lakes in the north-temperate zone, due to relatively transient episodes of high temperature and low oxygen, are predicted to increase more than fourfold by 2100 (Till et al. 2019). In order to predict and maybe counteract those catastrophic events, it is important to study how evolution has shaped extremophile organisms that can withstand both high temperatures and hypoxia (Tobler et al. 2018).

A system to study the joint effects of extreme temperatures and severe hypoxia is found in the sulfidic springs in Southern Mexico (Tobler et al. 2008b). Here, due to the discharge of H_2_S ground water from volcanic origin, dissolved oxygen is greatly reduced to severe hypoxia (often <1 mg/L O_2_, which can also be defined as anoxic but we would like to keep the term hypoxic as there are daily shifts and local variation in O_2_ levels, see results and (Tobler et al. 2006, Culumber et al. 2016, Lukas et al. 2021)). However, as these H_2_S-rich springs are of volcanic origin, water temperatures in the sulfidic habitat have been reported as being as high as 31.9°C (Jan 2006, (Tobler et al. 2008a)), which is well above temperatures found in adjacent clearwater river habitats.

Only two fish species, the sulfur molly, *Poecilia sulphuraria* (Álvarez 1948) and the widemouth gambusia, *Gambusia eurystoma* (Miller 1975) are regularly found in the sulfidic parts of the El Azufre River at the border between Tabasco and Chiapas. They are able to withstand the toxic effects of H_2_S and the H_2_S-related environmental hypoxia due to specialized adaptations at molecular (Pfenninger et al. 2014, Tobler et al. 2016, Passow et al. 2017, Tobler et al. 2017, Greenway et al. 2020, Kelley et al. 2021), morphological (Tobler and Hastings 2011, Tobler et al. 2011, Riesch et al. 2014, Passow et al. 2015, Greenway et al. 2016, Schulz-Mirbach et al. 2016), life-history (Riesch et al. 2010, Riesch et al. 2011a, Riesch et al. 2011b, Riesch et al. 2014, Jourdan et al. 2021) as well as behavioral levels (Plath et al. 2007, Tobler 2009, Lukas et al. 2021, Lukas et al. 2023).

Thermal tolerances (measured as CT_max_ and CT_min_) especially under hypoxic or anoxic conditions, however, have not been explored for these species although it is known that poeciliids in general have high thermal tolerance (40 to 43°C (Prodocimo and Freire 2001, Klerks and Blaha 2009, Bierbach et al. 2010, Culumber et al. 2015, Yanar et al. 2019, Nati et al. 2021). The majority of studies of resilience to environmental stressors have examined single stressors in isolation, whereas studies of the effects of interacting stressors are less common (Jackson et al. 2016). Thus, testing these extremophile fishes both for CT_max_ and CT_min_ under normoxic and hypoxic (e.g., natural anoxic) conditions will provide novel insights into evolutionary pathways allowing those fish to cope with multiple environmental stressors. Based on the ‘oxygen and capacity limitation of thermal tolerance’ hypothesis (OCLTT) we predicted lower thermal tolerances under anoxic conditions in our study species due to the fact that the aerobic metabolic demand cannot be met when under hypoxic conditions thus leading to a decrease in thermal tolerance (Pörtner 2001, McBryan et al. 2013, Jung et al. 2020, Earhart et al. 2022) (Pörtner et al. 2017, Jutfelt et al. 2018). Alternatively, there are several examples that show acclimation to hypoxia increases thermal tolerance in fishes (Burleson and Silva 2011a, Del Rio et al. 2021, Peruzza et al. 2021, Ern et al. 2023) and we could thus also predict that hypoxia might not affect thermal tolerances of our study species that are acclimated and even evolutionary adapted to severe hypoxia. Here, O_2_-independent anaerobic metabolism might be predominant which leaves these species unaffected in there thermal tolerances when tested under hypoxia.

Using laboratory based physiological assays on CT_max_ and CT_min_, we investigated how close these animals are to their critical temperatures encountered in their natural habitats. To this end, we estimated the heat tolerance margins of both species, e.g., the “safety gap” between highest temperatures possibly encountered in the natural habitat (as measured over several seasons with temperature loggers) and their physiological limits (Sunday et al. 2014, Desforges et al. 2023). Further we used the Thermal Death Time (TDT) framework (Ørsted et al. 2022) to calculate theoretical critical temperatures for both species.

## Material and Methods

### Study site

Our study system is located along a sulfidic spring complex near the city of Teapa in Tabasco, southern Mexico (the site is also known as Baños del Azufre’ site, 17°330 N, 93°000 W). Here a freshwater river is fed by the outflow of several groundwater springs which contain high levels of volcanic hydrogen sulfide (H_2_S, up to 990 μmol/l see: (Tobler et al. 2006, Culumber et al. 2016, Lukas et al. 2021)). This inflow creates a river stretch of approximately 2.5 km, which is well documented as an extreme aquatic environment, characterized not only by its high H_2_S content, but also the resulting low levels of dissolved oxygen and increased temperatures. In our study we concentrated on a river stretch of ca. 2 kilometers downstream the inflow of the first sulfur rich spring, in which the sulfur content is constantly above 170 μmol/l (Culumber et al. 2016).

#### Monitoring of water temperature regimes in the natural habitat

In order to establish the study site’s temperature regimes, we deployed HOBO temperature loggers (Onset Computer Corporation, Bourne, MA, USA) at five different locations along the river covering the main river channel as well as several springs and mixing areas (see map in **Figure 1** for logger locations). For the sulfidic part of the main river (Site 1, Figure S1), we were able to obtain a full year of hourly measurements (April 2018 to April 2019) while the other locations were measured during regular field trips (2018 to 2023, 1 week to 2 weeks periods, see Table S1 in the supplement).

**Figure 1:**
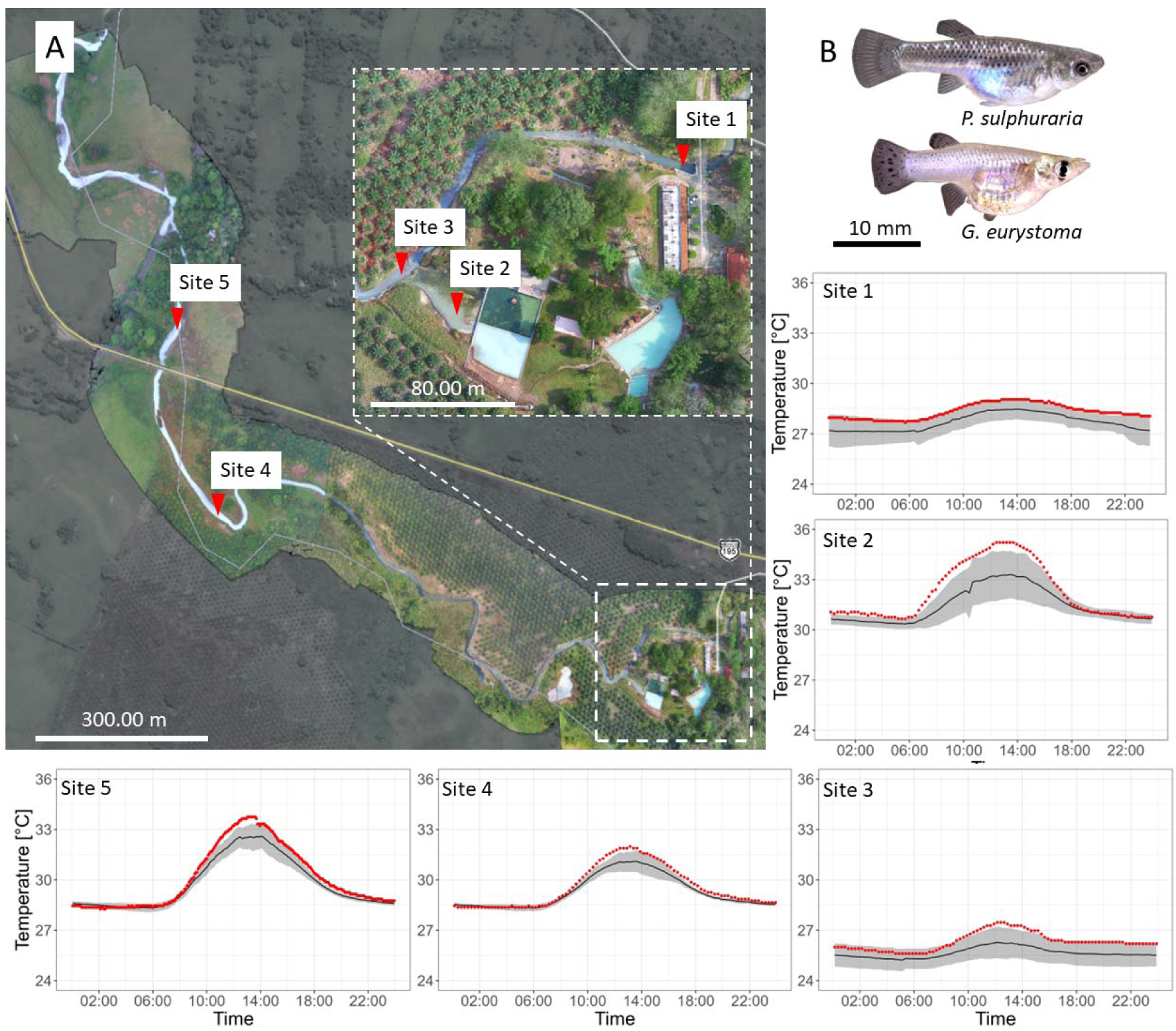
Location and temperature profiles of five study sites along a sulphidic spring system in Mexico and their inhabitants (*Poecilia sulphuraria & Gambusia eurystoma*). Temperatures were measured on a stretch of approximately one kilometer beginning shortly after the first sulfidic spring meets the main channel (A, site 1). Site 2 represents a location directly in front of a major sulfur spring, while sites 3 −5 represent various locations along the stream. Temperature profiles show the average daily temperature in °C ± s.d. (black line = mean over observation period; grey shaded area = s.d.; see Supplement: Table S1 for measurement periods from which means were derived for individual study sites). Red dots represent individual data points on the day in which the maximal temperature was observed. B: Female individuals of *Poecilia sulphuraria* and *Gambusia eurystoma*.

#### Determining upper and lower thermal tolerance limits of extremophile fishes

#### Sampling of study subjects and experimental protocols

##### May 23 experiment

CT_min_ and CT_max_ experiments under normoxic (100% O_2_ saturation) and severe hypoxic (0% saturation) conditions were conducted at the laboratories of DACBiol in May 2023. We collected widemouth gambusia (*Gambusia eurystoma*, N=60) as well as Sulfur mollies (*P. sulphuraria*, N=89) from the El Azufre river (site 5 in Figure 1, daily mean water temperature 18th May 5pm to 20th May 5pm: 29.9°C [range: 28.2°C to 35.2°C], season of the highest yearly water temperatures (see Figure 1, Supplement: Table S1, Figure S1)). Fish were collected with seines and dip nets the day of testing and kept at 30°C in cooler boxes containing a mixture of water from the collection sites and freshwater and were provided with aeration and filtration. Fish were visually matched for size prior to any experiment, however, *P. sulphuraria* were slightly larger than *G. eurystoma* (mean*_G.eurystoma_*: 17.8 mm, [range]: 14.9 mm - 26.1 mm standard length (SL); mean*_P.sulphuraria_*: 21.1 mm [15.3 mm– 32.6 mm]; *t*-test: *t*_145_=6.9, *p*<0.001).

The test apparatus for the CT_max_ experiments consisted of a 20-L glass tank with a circulating pump and an internal heating aggregate that were both placed inside a mesh cage to prevent fish from coming close. For the 100% O_2_ saturation treatment, an air-pump provided saturated oxygen concentrations throughout the tests. For the 0% O_2_ saturation treatment, we used 0.5 g/L Sodium-Sulfite to remove any oxygen from the water. This lack of oxygen represents normal conditions these fish experience at their site of collection (see Supplement: Figure S3).

We gently introduced on average 9 to 23 test fish from each species into the test tank and started increasing water temperatures after 20 minutes of habituation time. The temperature was raised at a constant rate of on average 0.42°C per minute. Water temperatures were recorded every minute with a measuring device (HACH HQ40D), and test subjects were monitored continuously. We removed the test fish separately once the test fish had turned its abdomen to the water surface and transferred the fish into an aerated 10-l tank at 30°C. All test fish regained motion control within a few minutes, and no mortality was associated with this experiment. After completion of a test trial, test fish were measured for SL using pictures on millimeter paper and ImageJ software.

For the CT_min_ experiment, we used the same apparatus and protocol but added ice cubes into the pump-holding mesh cage to ensure a constant decrease of temperature (decrease rate: 0.9°C per min).

We completed 1 trial for CT_max_ at 100% saturation, 3 trials for CT_max_ at 0% saturation, 1 trial for CT_min_ at 100% as well as 1 trial at 0% saturation.

In order to compare *C*_Tmax_ among the two species, we used a linear mixed model with species (*P. sulphuraria* and *G. eurystoma*) and O_2_-saturation level (100% or 0%) as well as their interaction term as fixed factors. We included trial as a random effect to take uncontrollable differences among the replicated trials into account.

CT_min_ values were compared in a linear model with species (*P. sulphuraria* and *G. eurystoma*) and O_2_-saturation level (100% or 0%) as well as their interaction term as fixed factors. Note that no random effect was included as there were no replicated trials per treatments in this experiment. Sample sizes per treatment can be found in Figures 2 and 3.

**Figure 2:**
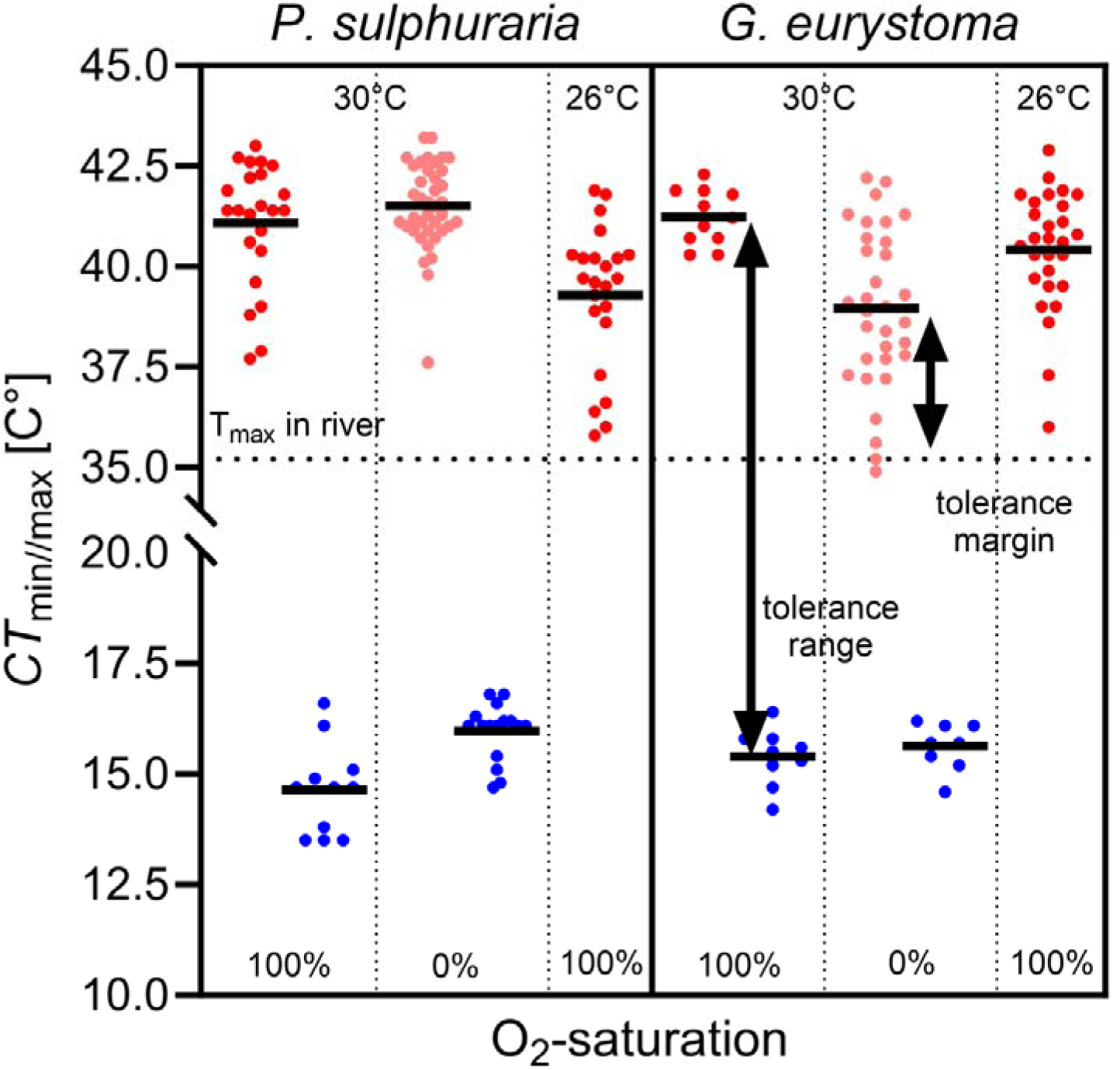
Critical temperatures (CT_max_ and CT_min_) of *P. sulphuraria* and *G. eurystoma* under 100% and 0% O_2_ saturation. For 100% saturation, CT_max_ of fish collected in May at 30°C and in February at 26°C water temperature is shown. Depicted are means along with all data points as well as the temperature range (CT_min_ to CT_max_) and the upper tolerance margin (T_max_ in habitat to CT_max_).

**Figure 3:**
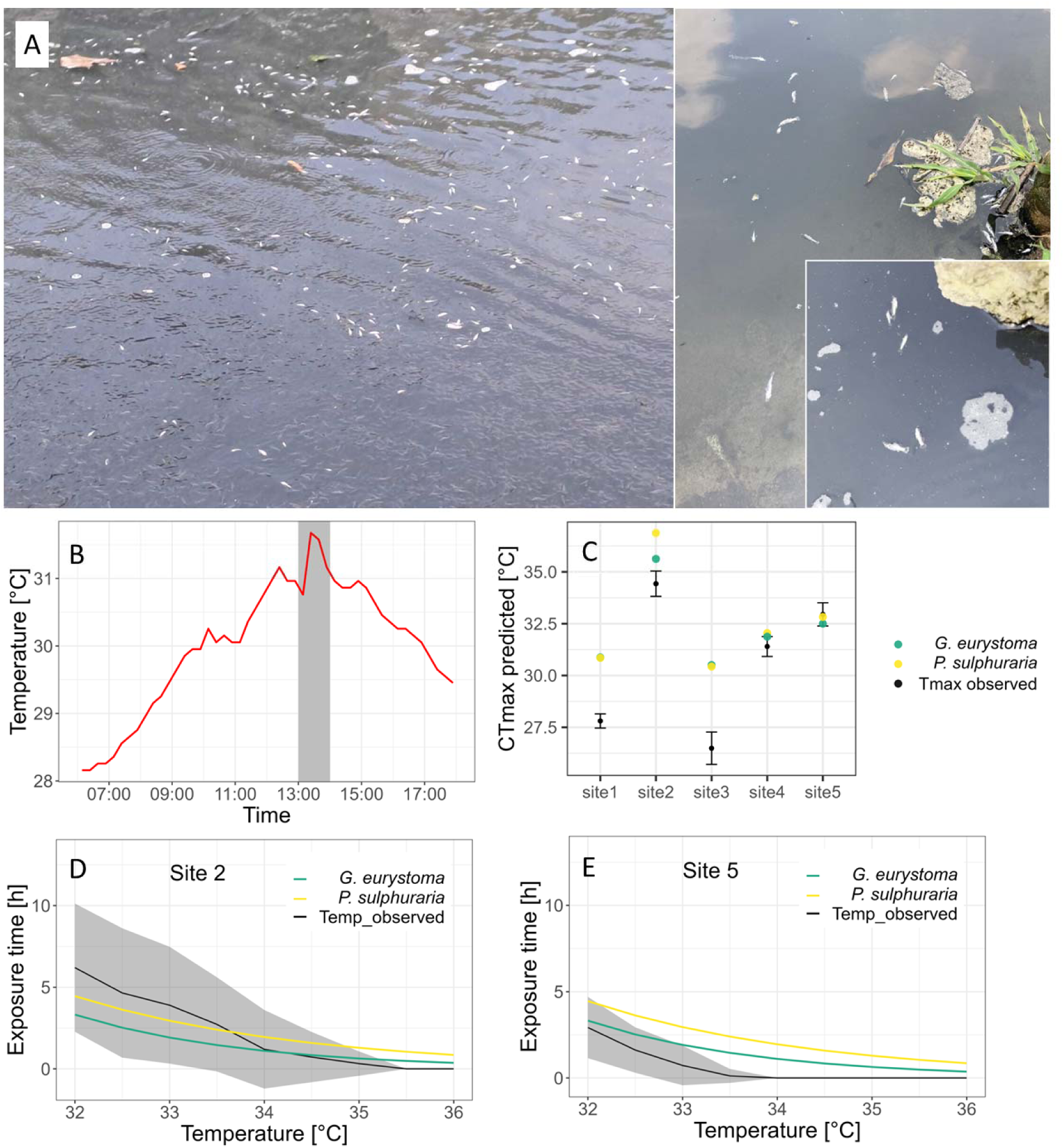
Fish mass mortality in habitat and predictions of TDT-modelling. (**A**) During our fieldwork in May 2023 we encountered sudden mass mortality of sulfur fish along the river. This coincided with the sudden release of an upstream reservoir filled with heated water, as indicated by an unusual increase in the temperature profile of site 4 on this day (**B**; grey area indicates time interval of mass mortality. (**C**) TDT model predictions for additional CT_max_ values in dynamic (heating) trials for *G. eurystoma* and *P. sulphuraria* under hypoxia with site-specific daily temperature increase as simulated heating rate (for site specific heating rate see Supplement: Table S3). Colored dots represent TDT model predictions, black dots with error bars indicates the mean site-specific daily temperature maximum with standard deviation. (**D;E**) Predicted tolerable exposure time to temperatures between 32°C and 36°C for *G. eurystoma* and *P. sulphuraria* under hypoxia and the mean time these temperatures were observed at the respective field sites. Colored lines represent TDT model predictions; black line represents the mean time interval certain temperatures were observed per day with grey shaded area representing standard deviation. The temperature range of interest was only measured in two field sites more than once (site 2; **D** and site 5; **E,** Supplement: Table S4).

To calculate the broad sense temperature range (breath) of both species, we subtracted treatment-specific average CT_min_ values from CT_max_ values.

##### February 23 experiment

In order to see how acclimation temperature may affect upper thermal limits, we further present experiments on CT_max_ under normoxic conditions and lowest river temperatures of the year (February 23) which we conducted in a field laboratory at CIIEA Teapa. Fish were collected at the same site and methods as described above (site 5, daily mean water temperature during sampling from 3:30 pm on 2nd February to 8am on 5th February: 25.8°C [range: 20.71 to 28.95]). Fish were kept one day before testing at 24.0°C in cooler boxes containing a mixture of water from the collection sites and freshwater and were provided with aeration and filtration. Fish were visually matched for size prior to any experiment (mean*_G.eurystoma_*: [range]: 20.4 [13.8–24.2] mm standard length (SL); mean*_P.sulphuraria_*: 20.8 [17.9–27.0] mm; unpaired *t*-test: *t*_33_=0.45, *p*=0.65).

The experimental design was identical to the one described above for the normoxic CT_max_ trials with the exception that fish were transferred into 24°C water after the completion of the experiment.

We completed 4 trials for CT_max_ at 100% saturation and compared CT_max_ among the two species using a linear mixed model with species (*P. sulphuraria* and *G. eurystoma*) as fixed factor and included trial as a random effect to take uncontrollable differences among the replicated trials into account.

### Calculating the physiological heating tolerance margins and theoretical critical temperatures

Lastly, we wanted to investigate if the temperatures observed in their habitat confront our study species with their actual thermoregulatory limits. Thus, we first calculated each species heating tolerance margins. We used the definition of heating tolerance margins given as the difference between the maximum water temperatures fishes experienced during the sampling period and their individual CT_max_ (CT_max_-T_max_habitat_).

Second, we applied a framework for a thermal death time (TDT) model and the respective R script developed and supplied in (Jørgensen et al. 2021, Ørsted et al. 2022). This framework allows for the prediction of tolerable temperatures at a given exposure time from data derived from dynamic heat experiments with a single heating rate, as carried out in this study. A crucial and highly sensible parameter in this approach is the heat sensitivity coefficient *z*, which must be estimated when predictions are done based on a single measurement (as our CT_max_ experiments in May were conducted with a constant heating rate, replicates in different treatment groups must be treated as a single measurement per group). To obtain a credible, yet conservative estimate for *z* we therefore used data from our February experiment. As heating rates between trials in February varied slightly (range 0.32 – 0.46°C/min), we used the observed CT_max_ in the individual trials in combination with their individual heating rates as data points to simulate the outcome of additional CT_max_ experiments (simulated heating rates: 0.2; 0.4, 0.5°C/min). These simulations supplied us with a predicted *z* value for *P. sulphuraria* and *G. eurystoma* respectively. We then used these values to predict CT_max_ for heating rates fish encounter in their habitat during the day and the duration of which temperatures in a certain range (32 – 36°C; observed water temperature maxima at our field sites) are tolerable for both species under anoxic conditions. For both simulations we used data based on the measurements we acquired in May. We are aware that model predictions based on such limited data conditions need to be interpreted with care, which is the reason why sensitivity coefficients for our predictions were calculated with fish acclimated to considerably lower temperatures (26°C). Further is the temperature range of interest in our model rather narrow and more importantly oriented towards the temperature range fish encountered in our experiments and in situ in their habitat (for a more detailed evaluation and complete model parameters see Supplement: Table S2, Figure S4, Figure S5).

#### Ethical and data statement

No fish died during our experimentation and fish were included in the stocks at DACBiol after the experiments were completed. The animal study was reviewed and approved by the Mexican “Comisión Nacional de Acuacultura y Pesca” (CONAPESCA; DGOPA.09004.041111.3088, PRMN/DGOPA-003/2014, PRMN/DGOPA-009/2015, and PRMN/DGOPA-012/2017). All data can be found in the supplement. Analyses were performed using SPSS 25 (IBM) and R (v4.2.2, https://cran.r-project.org/).

## Results

### Thermal regime of the El Azufre river

The clearwater river stretch merges with the first sulfide rich spring water at height of the Hacienda Los Azufres and thus temperature rises from daily averages of about 25°C to 28.8°C in the now sulfidic and hypoxic downstream stretch (site 1; see Figure 1). Water temperatures then increase along the river due to a widened river bed and several other inlets of sulfidic springs. In some adjacent pools that are shallow and with no contact to freshwater sources, water temperatures can increase to daily highs up to 35.2°C in the dry season (site 2; see Figure 1). Observed heating rates (measured from sunrise until the daily temperature maximum was reached) ranged from 0.003 °C/minute to 0.066 °C/minute and were therefore substantially lower than heating rates employed in our CT_max_ trials. Due to an release of hot sulfidic water from a pool used as a recreational swimming area into the main river, a sudden rise of water temperature of 1°C was detected 500 m downstream the release spot on 22th of May 2023. The regime of increased water temperatures lasted for one hour (see **Figure 3B**) and we observed thousands of dead and dying fish being flushed downstream (see **Figure 3A**, Supplement: video S1).

### CT_max_ under normoxic and severe hypoxic (anoxic) conditions

Sulfur mollies and widemouth gambusia differed in their upper thermal limits in relation to the water’s oxygen saturation (sig. interaction term of “species × saturation”, *F*_1,99_=15.7, *p*<0.001). While *G. eurystoma* had their critical maximal temperature at 100% O_2_ saturation with an average of 41.2°C [estimated marginal means; individual maximum recorded = 42.3°C] and showed a decrease in average CT_max_ to 39.0°C under hypoxic conditions (**Figure 2**), *P. sulphuraria* exhibited their highest CT_max_ at 0% O_2_ saturation with an average of 41.4°C (estimated marginal means; individual maximum recorded = 43.2°C) and a slightly lower CT_max_ under normoxic conditions (mean CT_max_ 41.1°C, **Figure 2**). Thus, both species only differed in CT_max_ values under hypoxic conditions (see **Figure 2**).

### CT_min_ under normoxic and severe hypoxic conditions

*P. sulphuraria* displayed their lowest critical minimum temperature at 0% O_2_ saturation while *G. eurystoma* did not differ in their CT_min_ in regard to O_2_ saturation (interaction term ‘species × saturation’: *F*_1,40_=5.4, *p*=0.025) but both species could tolerate lower temperatures when oxygen saturation was at 100% as compared to the 0% saturation treatment (oxygen: *F*_1,40_=11.1, *p*=0.002, **Figure 2**).

### Tolerance range (CT_max_-CT_min_)

Sulfur mollies showed their broadest tolerance range under normoxic conditions with a 26.4°C range and a slightly smaller range of 25.5°C at 0% saturation. Similarly, widemouth gambusia had their broadest range of 25.9°C at 100% saturation and a range of 23.3°C at 0% saturation.

### CT_max_ under normoxic conditions at 26°C collecting water temperature

Widemouth gambusia collected at a water temperature of 26°C in February 23 had a significantly higher CT_max_ under normoxic conditions (40.4°C) as compared to sulfur mollies with 39.2°C (*F*_1,46_=6.4, *p*=0.014; individual maxima: gambusia: 42.9°C, molly: 41.9°C, see **Figure 2**). Please note that average CT_max_ values at 100% O_2_ saturation for fish collected at 30°C water temperature (see above) were 1°C higher for widemouth gambusia and 2°C higher for sulfur mollies (see **Figure 2**).

### Thermal tolerance margins

As maximum water temperatures during the February sampling were at 28.95°C and 35°C during the May sampling, the thermal tolerance margins for both species were at 10.3°C and 11.5°C in February under normoxic conditions and under more natural anoxic conditions at 6.4°C and 3.9°C in May. In May, however, the range of individually recorded CT_max_ values for widemouth gambusia already overlapped with the maximum water temperatures (**Figure 2**).

### TDT modelling

Based on our February trials we obtained heat sensitivity coefficients *z* of 4.19 for widemouth gambusia and 5.58 for sulfur mollies respectively. Simulated heating trials based on our May data showed predicted CT_max_ values above the actual measured temperature maxima in all field sites except site 5, in which the mean measured temperature maximum was slightly higher than the predicted maximal temperature for both species given the observed heating rate (**Figure 3**). Water temperatures in our region of interest (32°C – 36°C) were repeatedly measured in site 2 and site 5. In site 5 the mean daily exposure times in that range were always below maximal tolerable exposure times predicted by the TDT model. At site 2, however, mean daily exposure times for temperatures in the range of 32°C – 34°C surpassed the predicted tolerable exposure time for both species substantially (**Figure 3**).

## Discussion

Using wild-caught fish that were acclimated to their H_2_S-rich, and at least during the daytime anoxic water, we found Sulfur mollies (*P. sulphuraria*) to show highest thermal resistance with CT_max_ values in several individuals exceeding 43°C under anoxic test conditions (O_2_ at 0% saturation) while widemouth gambusia (*G. eurystoma*) had their highest thermal limits under normoxic conditions and on average one degree Celsius lower than mollies under anoxia. For lowest tolerated temperatures, species differences became apparent only under normoxic conditions as sulfur mollies tolerated temperatures on average down to 14.5°C while widemouth gambusia had their lowest tolerated temperatures at 15.5°C. As environmental thermal regimes for multiple sites and years are available, we were further able to provide actual ecological context to our results. Thus, both fish species are confronted with temperature peaks (= environmental T_max_), only a few degrees Celsius below their critical temperature in their habitat. This is pointing towards a “life on the edge” as well as a low ability to withstand further temperature extremes that might become more common in future due to global change, although we found these fish to increase their CT_max_ to some degree when acclimated to warmer water. A recorded mass killing due to an sudden release of hot water into the main river that led to a sudden increase of water temperatures of 1°C further exemplifies that the fishes in this system, although there are highly adapted, live at the very upper edge of their physiological limits.

Sulfur adapted fishes in the El Azufre system face a threefold suit of abiotic stressors with H_2_S, severe hypoxia and elevated temperatures. Interestingly, both fish species differ in their tolerance towards high and low temperatures under anoxic conditions. Without O_2_ in the water, which represents normal conditions during the day in this system (see SI, (Culumber et al. 2016, Lukas et al. 2021), sulfur mollies showed highest thermal resistance with CT_max_ values in several individuals exceeding 43°C. Although these fish had been acclimated to anoxic conditions as they were wild-caught and tested timely after capture, the fact that Sulfur mollies’ thermal maximum was not affected by anoxic conditions at all is astonishing and unprecedented to our knowledge.

*G. eurystoma* on the other hand showed a pattern known from other tropical fish, mainly that CT_max_ is reduced under hypoxic conditions. In a study on zebra fish, larvae exposed to hypoxic conditions showed lower CT_max_ then those exposed to normoxic while those exposed to hyperoxic conditions showed highest CT_max_ values (Andreassen et al. 2022). Still, the ability to maintain CT_max_ values under anoxic conditions that were close (only 1°C below) to those under normoxic conditions is also an extraordinary adaptation in this extremophile species.

It is known that *P*. sulphuraria and to a lesser extent also *G. eurystoma* exhibit differential expression of and positive selection on oxygen transport genes (Barts et al. 2018, Greenway et al. 2020) and up-regulation of genes associated with anaerobic ATP production (Kelley et al. 2016). Further, sulfid-adapted fishes show increased head size along with increased gill surface areas (Tobler et al., 2011) that is both correlated with ventilation efficiency (Camarillo et al. 2020). Thus, oxygen uptake through ASR behavior (Lukas et al. 2021) could be still sufficient under aquatic anoxia for these species to satisfy their metabolic demand even at high temperatures. At the moment, we do not know to what extent acclimatization (along with epigenetic changes, see (Kelley et al. 2021)) to anoxic or hypoxic conditions plays a role in that tolerance to high temperatures as found in other fish. For example, hypoxia acclimation of channel catfish (*Ictalurus punctatus*) was found to increase the cardiovascular ability to withstand an acute temperature increase and thus led to higher CT_max_ (Burleson and Silva 2011b). Similarly, also so-called acquired cross-tolerances are a possible explanation through exposure to one stressor (hypoxia) can increase tolerance towards another (temperature in our case (Rodgers and Gomez Isaza 2021)). For example, in Chinook salmon, *Oncorhynchus tshawytscha*, heat tolerance was improved by short term exposure to high salinity and air which exemplifies that some forms of stress can heighten acute heat tolerance in ectotherms (Rodgers and Gomez Isaza 2022).

In order to evaluate how stressor combinations and acclimatization may effect tolerances in both species towards H_2_S, hypoxia and elevated temperature, future research with simulated stressor environments are needed although this is technically highly demanding. Furthermore, this system seems to be suited to disentangle molecular and physiological mechanisms underlying heat stress, hypoxia and H_2_S tolerances through sophisticated -omics approaches (see for example (Payne et al. 2022)).

While Sulfur mollies and widemouth gambusia showed Ct_max_ values at the very upper end of those reported for tropical fishes, they do not overshoot or have exaggerate tolerances per se (see values from other poeciliids in (Bierbach et al. 2010, Nati et al. 2021)). Given their thermally extreme habitat, we found that they already live at temperatures close to their CT_max_ which means they have small thermal safety margins (Comte and Olden 2017). This assumption is further supported by the predictions obtained via the application of a TDT model framework. The simulation of additional heating trials based on our experimental results and heating rates observed in the river showed that both species are regularly confronted with their CT_max_ under natural conditions (i.e., site specific observed heating rates) at several locations throughout their habitat. Furthermore, did observed time intervals of elevated temperatures exceed the predicted duration these temperatures were theoretically tolerable for both species. Even though our TDT calculations are based on limited data, the results do further highlight the constant thermoregulatory challenges both Sulfur-adapted fish species face in their habitat. Predicted values the time fish can tolerate certain temperatures show that increased intervals of temperatures in the range of 32 °C – 34°C could present a greater challenge for our study species than shorter periods of extreme temperature maxima. As tropical ectotherms are generally expected to be vulnerable to human-induced climate change for various reasons (Sunday et al. 2014, Desforges et al. 2023), it is reasonable to assume that an increase of 2°C water temperature in our sulfur riverine system may render substantial portions uninhabitable for *P. sulphuraria* and *G. eurystoma.* The observation of the described event of mass mortality did further highlight the extreme fragility of the studied ecosystem. While we cannot completely rule out a possible role of increased H_2_S in the event (measures not available), a measurable temperature increase of 1°C for 1 hour at a field site 800 m downstream suggests a strong role of elevated temperatures at the origin of the event (i.e., where non-flowing heated water was released). We therefore assume that fish further upstream were confronted with a temperature increase that possibly exceeded the maximum duration fish could tolerate these temperatures. In addition, the flashflood like nature of the event might have prevented migration in more thermally favorable micro-habitats along the river and therefore can be seen exemplarily for the effects of a constant rise in water temperature and its effect on both species. This comes with alarming implications as both species are listed as globally endangered (*G. eurystoma*: CR, *P. sulphuraria*: EN) by the IUCN due to their narrow natural distribution (IUCN 2022).

## Conclusion

In supplying experimental data for CT_max_ under normoxic and hypoxic conditions in two extremophile fish species and the direct comparison to temperature measurements in their habitat we could show that sulfur mollies and wide mouth gambusia are regularly experiencing *in-situ* water temperatures close to their thermal limits. We conclude that, while the role of physiological acclimatization and evolutionary adaption capability are still a challenging aspect in predicting the influence of global warming on ectotherm species, our study system might represent an example in which a minimal increase in sustained water temperature is sufficient to threaten the existence of two endemic and already endangered species. Thus, more research in how multiple environmental stressors interact and alter the physiological performance in these fishes may yield the potential to understand and predict the challenges of anthropogenic influences on further ecosystems.

## Acknowledgement

We are grateful to the director and staff at the CIIEA Centro de Investigación e Innovación para la Enseñanza y el Aprendizaje field station for hosting our multiple research stays. We thank Carla Vollmoeller, Charlotte Steven and Anna Helmke for assistance during the fieldwork and the owner of Antiguo Jacalito for his generosity.

## Author contribution

LAR, NHR, KP, JK and DB conceived the study. KP, NHR, DB, JL, CV and YS measured water parameters. KP, NHR, DB, AJL, EJJ and LAR performed the experiments. DB, KP and NHR analyzed the data. KP and DB wrote the first draft of the manuscript which was commended, adjusted and finally approved by all authors.

