## Supplemental material for "Increased thermal tolerance under anoxic conditions in an extremophile fish from hot sulfur springs in Mexico"

**SUPPLEMENTARY INFORMATION**

**Table S1: Location, temperature measurement periods, maximal, minimal, and mean measured temperature for five field sites in the ‘Azufre’ sulfur riverine system**. Note the two measurement intervals at site 5. Temperature data for February is included as fish for our trials in February were caught under this temperature regime.

| **Site #** | **Location (lon,lat)** | **start** | **end** | **max (C°)** | **min (C°)** | **mean (C°)** |
| --- | --- | --- | --- | --- | --- | --- |
| 1 | 17.552779, -92.997249 | 05/04/2018 | 22/04/2018 | 29.1 | 24.4 | 27.4 |
| 2 | 17.552211, -92.998429 | 17/03/2022 | 26/03/2022 | 35.2 | 29.4 | 31.5 |
| 3 | 17.552265, -92.998695 | 17/03/2022 | 26/03/2022 | 27.5 | 24.2 | 25.6 |
| 4 | 17.554398, -93.004935 | 14/05/2023 | 24/05/2023 | 31.9 | 28.0 | 29.3 |
| 5 | 17.554398, -93.004935 | 01/02/2023 | 04/02/2023 | 29.1 | 23.9 | 26.6 |
| 5 | 17.557412, -93.006040 | 14/05/2023 | 24/05/2023 | 33.7 | 27.9 | 29.8 |

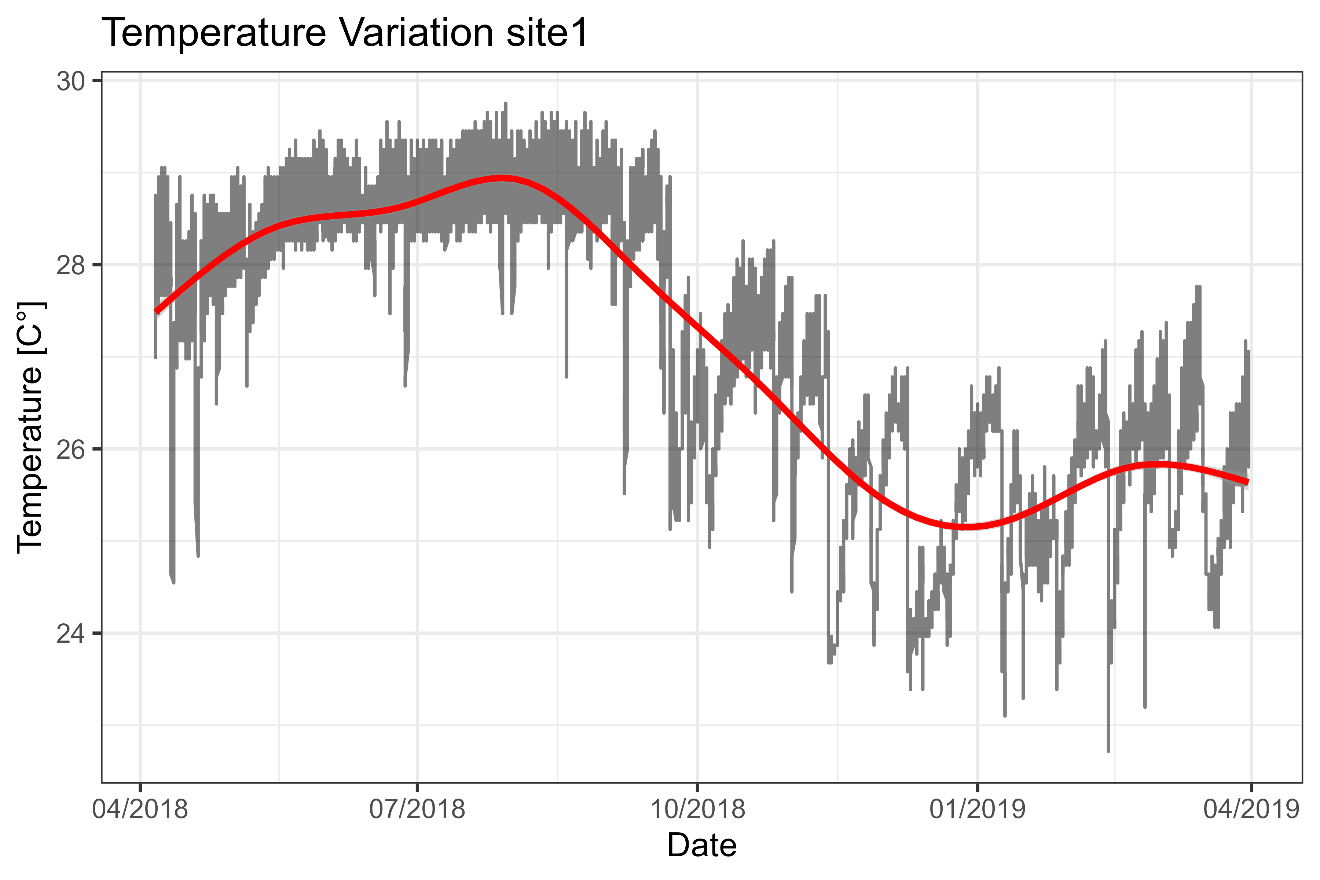

**Figure S1**: **Water temperature profile of a yearlong measurement from 04/2018 – 04/2019 at site 1**. Red line indicates average temperature. Grey line indicates hourly temperature measurements. Please note the overlap in measurement period with values shown in table S1. The values in table S1 were acquired with a different temperature logger with a 10-minute measurement interval.

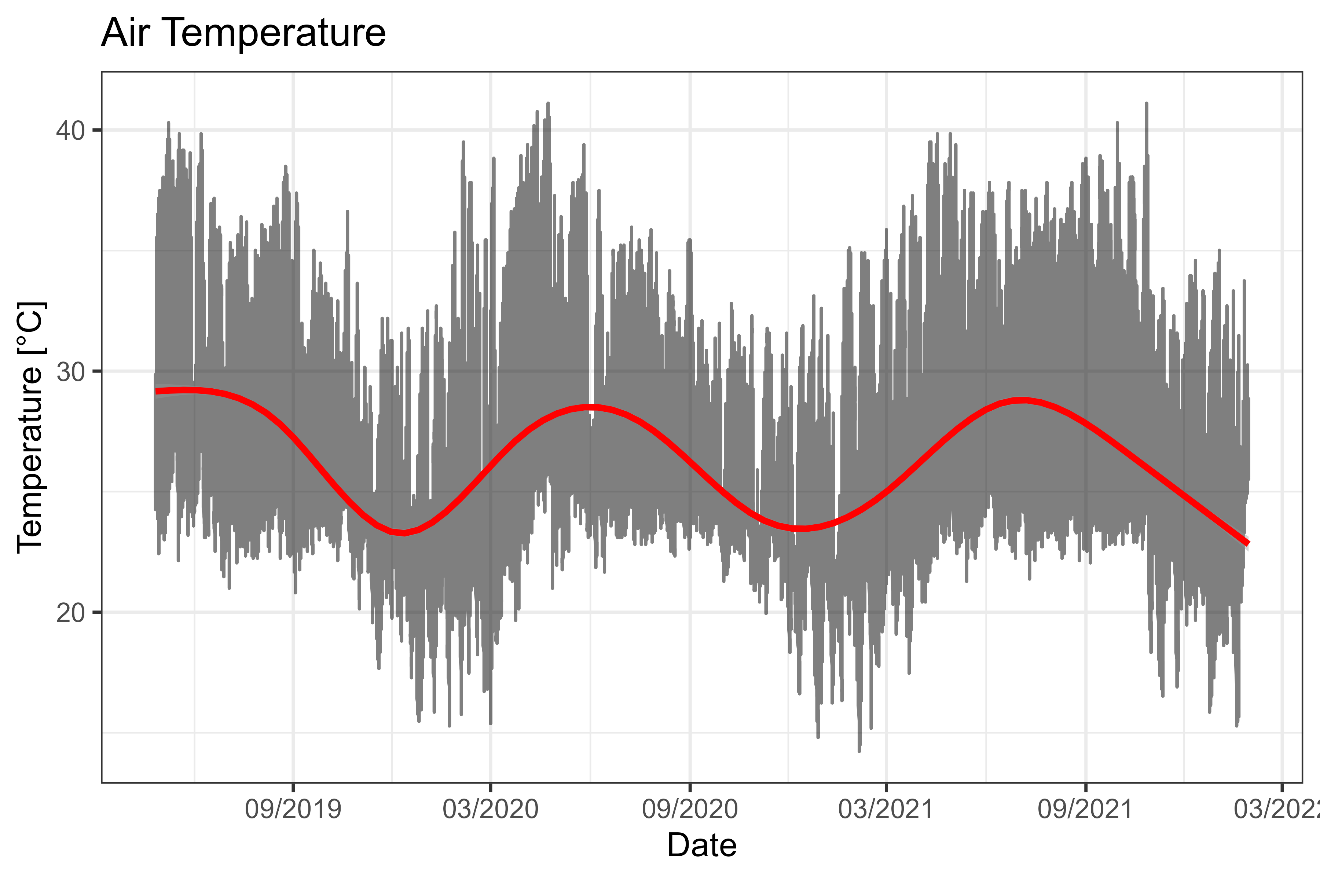

**Figure S2**: **Temperature profile of air temperature measured near field sites 1-3 from 4/2019 – 3/2022.** Red line indicates average temperature. Grey line indicates individual hourly temperature measurements.

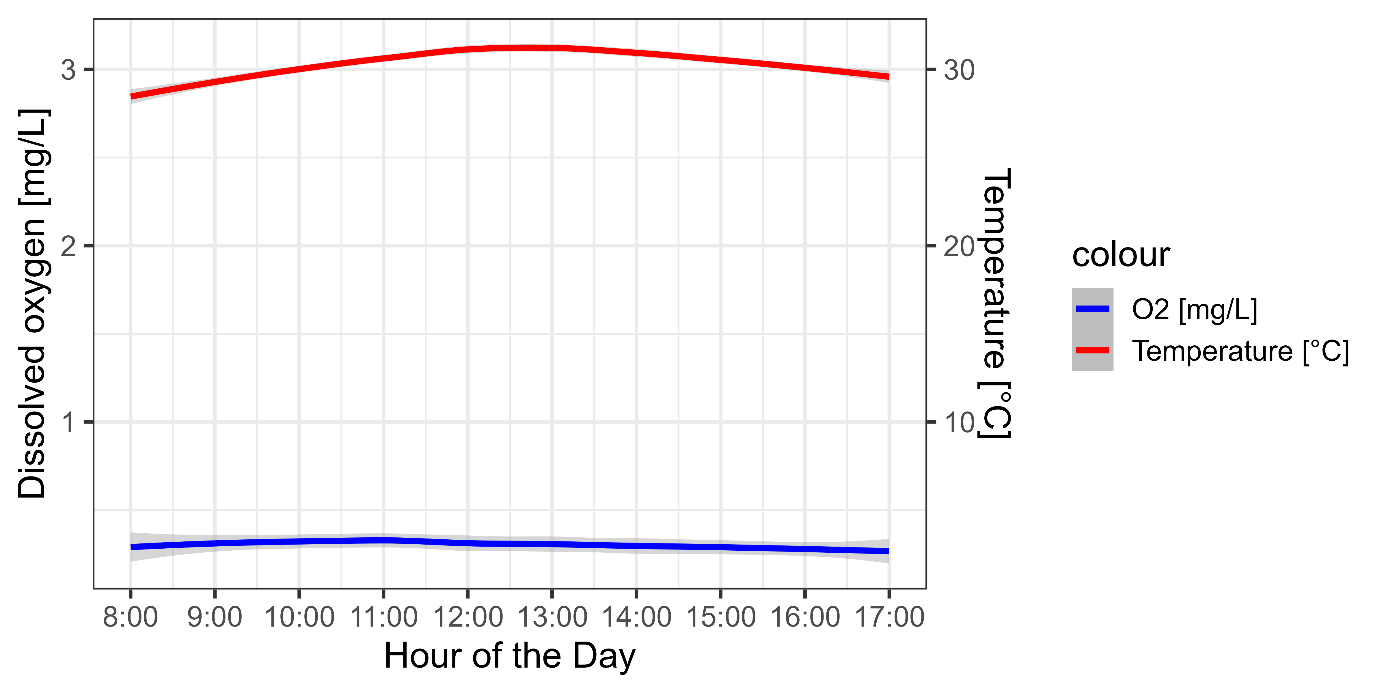

**Figure S3: Dissolved oxygen (blue, left axis) and temperature (red, right axis) during the day on our field sites in May 2023**. Measurements were taken during the day (8:00 – 17:00) twice for every field site measured in May 2023 (site 4 & 5). Grey shaded areas indicate 95CI.

**TDT Model framework**

The framework supplied by (Jørgensen et al., 2021) allows for predictions based on a single measurement. As mentioned in the main text we are aware of the pitfalls that come with specific model predictions and especially the estimation of *z*. We are, however, confident that our predictions have a sufficient level of reliability for the following reasons: Firstly, our *z-values* are derived from actual experimental data (February 2023) on both species. While the range in heating rates we used to derive the values may be narrow, they fall well into the upper limit of z-values reported in the SI of Jørgensen et al. (2021, range provided: z = 1.4 – 5.8), as you would expect from species adapted to high temperatures. Secondly, we restricted our predictions to both temperature and time intervals close to the testing conditions in our experimental setting. This is crucial to avoid extrapolation uncertainties induced by predictions outside the respective thermal or temporal domain.

**Table S2: TDT parameters used to predict additional CT_max_ values and maximum tolerable temperature**. Note that only predictions for no oxygen treatment is shown in the manuscript as this treatment mirrors the conditions fish encounter in their habitat.

| *Treatment* | *z* | *T_0_ (C°)* | *T_C_ (C°)* | *dCT_max_ (C°)* | *Ramprate (C° min^-1^)* |
| --- | --- | --- | --- | --- | --- |
| P. sulphuraria oxygen | 5.58 | 30 | 28 | 41.4 | 0.45 |
| P. sulphuraria no oxygen | 5.58 | 30 | 28 | 39.5 | 0.44 |
| G. eurystoma oxygen | 4.19 | 30 | 28 | 40.6 | 0.45 |
| G. eurystoma no oxygen | 4.19 | 30 | 28 | 39.0 | 0.44 |

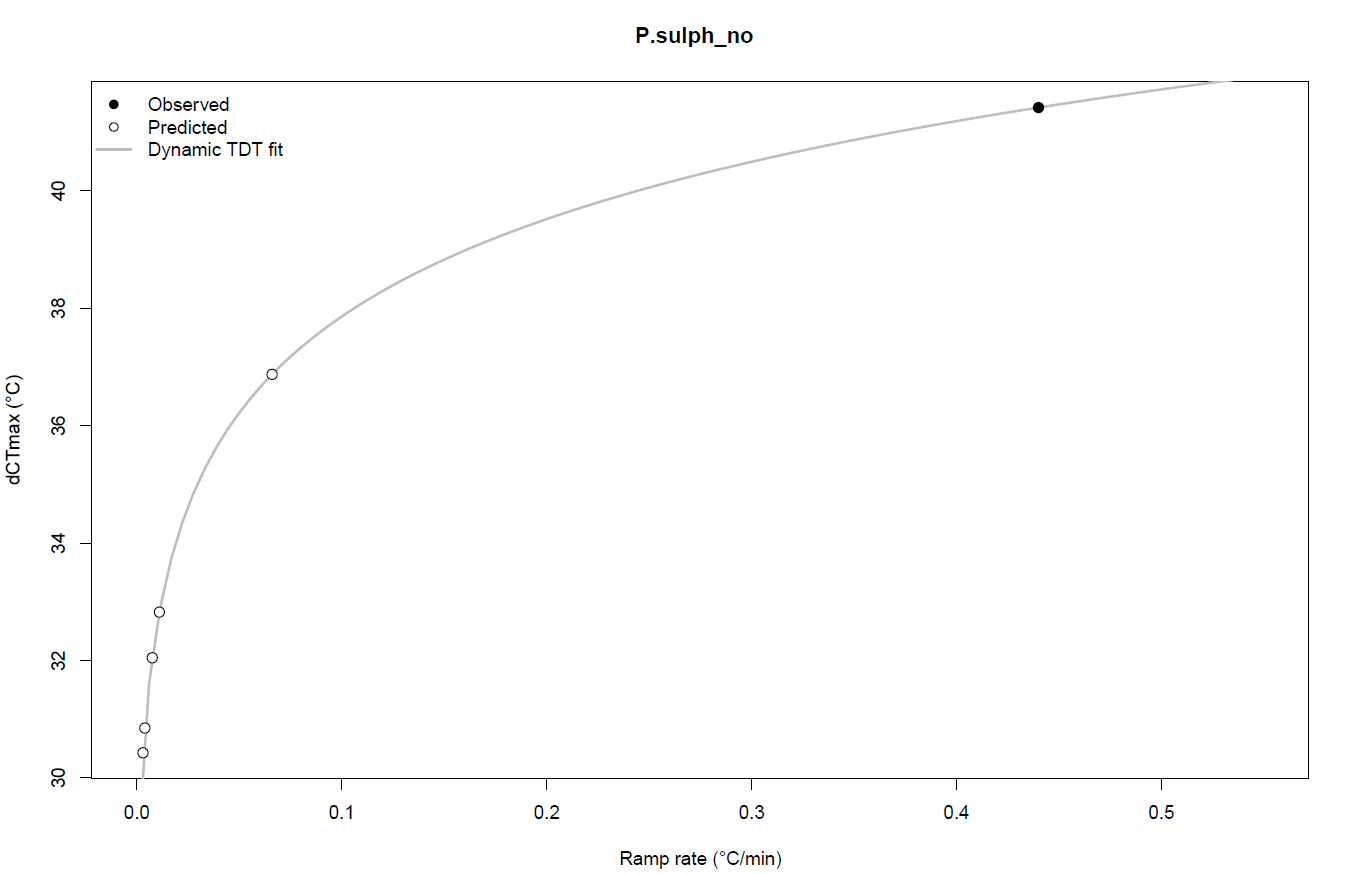

**Figure S4: TDT curve for *P. sulphuraria* under hypoxia predicted for observed heating rates in habitat (white circles) and the results of heating trials in May 2023 (black circle).** For further model parameters see table S4.

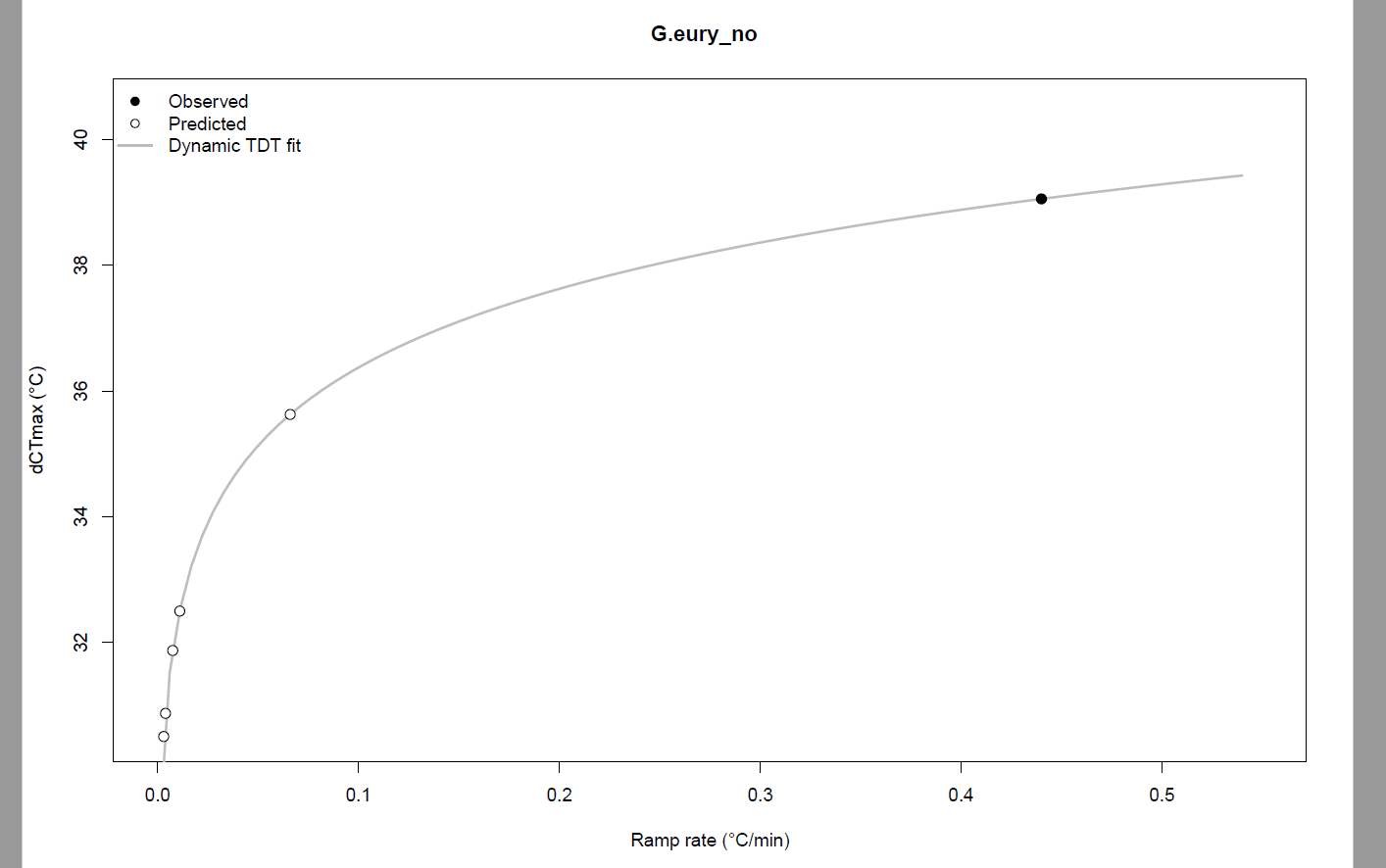

**Figure S5: TDT curve for *G. eurystoma* under hypoxia predicted for observed heating rates in habitat (white circles) and the results of heating trials in May 2023 (black circle).** For further model parameters see table S4.

**Table S3: Predicted CT_max_ for simulated trials based on observed heating rates**. Heating rates for individual field sites were calculated as mean daily temperature increase over time from sunrise until the daily temperature maximum was reached. Additional CT_max_ predictions are based on these heating rates and can be compared to the maximum temperature (T_max_) measured at specific sites.

|  |  | **CT_max_ predicted [°C]** | |  |
| --- | --- | --- | --- | --- |
| **Site** | **Observed heating rate [°C/min.]** | ***G. eurystoma*** | ***P. sulphuraria*** | **T_max_**  **observed [°C]** |
| 1 | 0.0039 | 30.9 | 30.9 | 29.1 |
| 2 | 0.066 | 36.9 | 35.69 | 35.2 |
| 3 | 0.003 | 30.4 | 30.59 | 27.5 |
| 4 | 0.0075 | 32.0 | 31.9 | 31.9 |
| 5 | 0.011 | 32.8 | 32.5 | 33.7 |
| **CT_max_ trial May** | **0.44** | **41.4** | **39.0** |  |

**Table S4: Daily exposure to heat levels above certain temperatures *P. sulphuraria* and G*. eurystoma* encounter in their habitat and predictions for tolerable exposure times at those temperatures.** Shown are TDT model predictions based on the results of oxygen deprived CT_max_ experiments from May 2023, as this reflects conditions fish face in their habitat. Site 2 and Site 5 were the only field sites in which temperatures above 32 °C were observed more than once in the measurement period.

| **Temperature (°C)** | **Mean exposure time per day ± s.d. in habitat (minutes)** | | **Predicted time of exposure tolerated (minutes)** | |
| --- | --- | --- | --- | --- |
|  | **Site 2** | **Site 5** | ***G. eurystoma*** | ***P. sulphuraria*** |
| 32 | 372.0 ± 235.6 | 175.5 ± 106.5 | 199.3 | 267.3 |
| 32.5 | 279.0 ± 237.7 | 97.3 ± 78.9 | 151.4 | 217.5 |
| 33 | 234.0 ± 214.6 | 43.6 ± 68.9 | 115.0 | 176.9 |
| 33.5 | 163.5 ± 173.0 | 7.3 ± 24.1 | 87.4 | 143.9 |
| 34 | 72.0 ± 144.4 | 0.0 ± 0.0 | 66.4 | 117.1 |
| 34.5 | 43.5 ± 92.3 | 0.0 ± 0.0 | 50.4 | 95.3 |
| 35 | 19.5 ± 44.7 | 0.0 ± 0.0 | 38.3 | 77.5 |
| 35.5 | 0.0 ± 0.0 | 0.0 ± 0.0 | 29.1 | 63.1 |
| 36 | 0.0 ± 0.0 | 0.0 ± 0.0 | 22.1 | 51.3 |
